# DOK1 and DOK2 regulate CD8^+^ T cell signaling and memory formation without affecting tumor cell killing

**DOI:** 10.1101/2021.12.17.473111

**Authors:** V. Laletin, P.-L. Bernard, C. Montersino, Y. Yamanashi, D. Olive, R. Castellano, G. Guittard, J. A. Nunès

## Abstract

Targeting intracellular inhibiting proteins is a promising strategy to improve CD8^+^ T cell anti-tumor efficacy. DOK1 and DOK2 are CD8^+^ T cell inhibitory proteins that are targeted in this study in order to improve the activation and cytotoxic capacities of these cells. To evaluate the role of DOK-1 and DOK-2 depletion in physiology and effector function of T CD8^+^ lymphocyte and in cancer progression, a transgenic T cell receptor mouse model specific to melanoma antigen hgp100 (pmel-1 TCR Tg) was established.

Depletion of both *Dok1* and *Dok2* did not affect the development, proliferation, mortality, activation and cytotoxic function of naive CD8^+^ T cells. However, after an *in vitro* pre-stimulation *Dok1*/*Dok2* DKO CD8^+^ T cells had higher percentage of effector memory T cells and showed an increase in levels of pAKT and pERK upon TCR stimulation. Despite this improved TCR signaling, pre-stimulated *Dok1*/*Dok2* DKO CD8^+^ T cells did not show any increase in their activation or cytotoxicity capacities against melanoma cell line expressing hgp100 *in vitro*.

Altogether we demonstrate here a novel aspect of the negative regulation by DOK1 and DOK2 proteins in CD8^+^ T cells. In conclusion, DOK1 and DOK2 have an inhibitory role following long term T cell stimulations.

## INTRODUCTION

CD8^+^ T cells have a key role in tumor eradication through their capacity to specifically recognize tumor antigens and to secrete potent effector molecules for tumor cell killing such as IFNγ, TNFα and granzymes.^1–3^ The potential of these cells is a foundation for modern approaches of cancer immunotherapy. Several CD8^+^ T cell-based methods were proposed to fight cancers: CAR-T cells, TILs and vaccine-based approaches.^4–6^ Despite the potency of these methods the majority of patients do not respond. The reason underlying this phenomenon is associated with the negative regulatory tumor microenvironment, inhibitory ligands and diminished TCR signaling.^7–11^ Therefore, new methods to improve CD8^+^ T cell-based immunotherapies of cancer are required. One promising approach is to improve TCR signaling targeting intracellular inhibiting proteins.^12^

TCR signaling is crucial for T cell activation, differentiation and cytotoxicity. It is initiated by surface expressed T cell antigen receptor TCR, extended by complex step of intracellular signal transduction and potentiation that implicates numerous proteins.^13^ Protein tyrosine kinases, such as Lck, Fyn and ZAP-70 are involved in proximal to TCR signal transduction, whereas adaptor proteins LAT, Grb2 and SLP-76, intracellular signal transducers PI3K and RAS form a key signalosome for distal to TCR signal potentiation.^14,15^ Finally, these TCR encoding signals lead to activation of two main effector pathways in T cell activation, RAS/ ERK-1/2 and PI3K/ AKT signaling pathways that could be detected by the ERK-1/2 and AKT phosphorylation status. Upon TCR engagement and activation, naïve T cells undergo a maturation process that give rise to memory T cells. This maturation is accompanied with changes in TCR signal encoding, activation and expression of surface markers.^16,17^ Therefore, TCR signaling pathway may differ between naïve and memory, CD4^+^ and CD8^+^ T cells. ^18–22^

Previously it was shown that targeting an intracellular TCR-signaling inhibiting protein CISH can improve TCR activation and tumor clearing in tumor bearing mice model.^23,24^ Based on these studies, a clinical study invalidating CISH in CD8^+^ tumor infiltrating lymphocytes (TILs) via Crispr-CAS9 approach prior to adaptive cell transfer is ongoing (NCT04426669). Thus, a new concept of cancer immunotherapies to target intracellular inhibiting proteins is emerging. Here we investigated the targeting of intracellular TCR-signaling inhibiting proteins DOK1 and DOK2 to improve CD8^+^ T cell activation and cytotoxicity against tumors.

Initially identified as a tumor suppressor genes, DOK1 and DOK2 adaptor proteins are two members of DOK family proteins that are constitutively expressed in T cells.^25–28^ DOK1 and DOK2 expression increase upon T cell maturation.^29^ TCR engagement induces the phosphorylation of DOK1 and DOK2.^29–31^ DOK1 and DOK2 proteins are implicated in negative regulation of TCR signaling as their deficiency improve TCR-mediated cytokine production and proliferation in T-cell lines and in mouse CD4^+^ T lymphocytes.^29,30^ Upon tyrosine phosphorylation of DOK1/2, some DOK-interacting proteins such as RasGAP, SHIP and Csk proteins are involved in negative feedback loops for TCR signal transduction.^14,25,26,28–30,32,33^ After TCR engagement, the loss of DOK1 and DOK2 in CD4^+^ T cells increases both early phosphorylation events such as ZAP-70 and LAT and distal phosphorylation events as ERK-1/2 and AKT. ^29,30^ Very few data are available for the CD8^+^ T cell compartment. Previously, it has been reported that the CD3 ligation induces the DOK2 tyrosine phosphorylation in a human cytotoxic T cell clone.^30^ The analysis of CD8^+^ T cells in *in vivo* infection mouse model showed higher production of IFNγ, TNFα and granzyme B in DOK1 DOK2 KO mice, without affecting TCR signaling. ^31^

To further understand how DOK1 and DOK2 regulate CD8^+^ T cell activity and especially their cytotoxic function against cancer cells, we crossed *Dok1*/*Dok2* DKO mice with pmel-1 TCR transgenic mice.^23^ We then investigated the role of DOK1 and DOK2 in physiology, TCR signaling and activation of naïve and primed CD8^+^ T cells and cytotoxic capacity against tumor. Primed *Dok1*/*Dok2* deficient CD8^+^ T cells showed increased TCR signaling evaluated by ERK-1/2 and AKT phosphorylation that was not observed in naïve CD8^+^ T cells and acquired effector memory phenotype upon CD8^+^ T cell amplification. However, we detected no difference in activation, cytokine production or cytotoxicity against tumor cells.

We demonstrate, here, a novel aspect of the negative regulation by DOK1 and DOK2 proteins in CD8+ cells. Indeed, our results allow us to conclude that DOK1 and DOK2 have an inhibitory role following longer term stimulations. This mechanistic knowledge advances our understanding of T cell function and may lead to novel approaches that enable the development of enhanced *in-vitro* T cell strategies for cancer immunotherapy.

## MATERIALS AND METHODS

### Mice and cell lines

#### Mice

In brief, *Dok1 Dok2* KO mice (C57BL/6) were obtained from Yuji Yamanashi, University of Tokyo, Japan. Pmel-1 TCR transgenic mice, were a kind gift from Nicholas P. Restifo (NCI, Bethesda, USA).^34^ Mice were crossed to generate *Dok1 Dok2* KO Pmel-1 transgenic mouse strain. All mice were crossed, housed and genotyped according to the guidelines of Committee for Animal Experimentation of Marseille and in accordance with European Directive 2010/63/EU. The experimental protocol was approved by an Institutional Animal Care and Use Committee. Male and female mice were used between the ages of 6–12 weeks. Mice were bred and maintained under specific pathogen-free conditions at the Centre de Recherche en Cancérologie de Marseille (CRCM) animal facility.

#### Genotyping

Pmel-1 TCR genotyping was performed as described previously^23^ using following primers: 5Chr2pmel: 5’ CTT TAG ACC TCC GGC ACT GTT GC 3’; 3Chr2pmel: 5’ GCA AGT AGC AGT GTA TCA AAT ATG C 3’; 3PmelTCRb: 5’ GTA GCT TTG TAA GGC TGT GGA GAG 3’, with expected bands band sizes at 280 and 300 bp. Dok1 and Dok2 genotyping was performed using following primers: Dok1_F bisGAAATGACATCTTTCAGGCAGTTGAGGC; Dok1_R bis GAGTCTGTCAGCTTGGTTTTCAGTAACT; Dok2_F GTTCGCAGCCGTGTTATATGGAGAGTCT; Dok2_R GAAAGCCAACAGGCAGATGGCCTGTAT, with expected bands on 351 and 261 bp for Dok1 and Dok2 respectively.

#### Cell culture

B16 melanoma (H-2D^b^), a mouse melanoma, transduced retrovirally to express human glycoprotein 100 (hgp-100) with human residues at positions 25–27 was a kind gift from the team of Nicholas P. Restifo (NCI, Bethesda, USA).^35^ WT B16 melanoma cells and B16 expressing hgp-100 were grown in DMEM 20%FCS, 1%NEAA, and 1% Sodium butyrate.

### *In vitro* CD8^+^ T cell expansion

Naïve CD8^+^ T lymphocytes were isolated from splenocytes by magnetic bead negative selection per the manufacturer’s protocol (Invitrogen). CD8^+^ T cell expansion was performed in RPMI medium 10%FCS, 50μM b-mercaptoethanol. Splenocytes were cultured in presence of hgp100_25–33_ peptide, KVPRNQDWL (AnaSpec, CliniSciences) at 100ng/mL and IL-2 (100IU/mL) or expanded by plate-bound anti-CD3 mAb at 2μg/mL (BD Pharmingen) and soluble anti-CD28 mAb at 1μg/mL (BD Pharmingen) and IL-2 (100IU/ml) for 3 days. This was followed by 2 days of IL-2 (100IU/ml) maintenance.

### Flow cytometry

For cell phenotyping a single cell suspension was prepared. Red blood cells lysis was performed if necessary, using 1X ACK lysis buffer (Gibco). Extracellular staining was performed for 30 minutes at 4°C. When necessary, intracellular staining was performed by use of the FoxP3/Transcription Factor Staining Buffer Set (eBioscience) according to the manufacturer’s instructions. Used antibodies for CD8^+^ T cell phenotyping: CD4-Percp5,5 (BioLegend, #100539), CD3-PE-Cy7 (eBioscience, #25-0031-82), CD8-APC-EF780 (Invitrogen #47-0081-82), CD62L-APC (Invitrogen, #17-0621-81), CD44-FITC (BD Pharmingen, #561859), CD62L-eF450 (Invitrogen, #48-0621-82), CD44-AF700 (Invitrogen #56-0441-82). For phosphoflow experiments cells were immediately fixed by FoxP3/Transcription Factor Staining Buffer Set (eBioscience) for 10 minutes at 37°C after stimulation and labeled by pErk-AF488 (Cell Signaling, #13214) and pS6-APC (Cell Signaling, #14733) antibodies. Dead cell exclusion was done by LIVE/DEA Fixable Aqua Stain (Thermo Fisher Scientific, L34957). All data were acquired on LSRII, Fortessa, (BD Biosciences) and analyzed with FlowJo software (Tree Star, Ashland, OR, USA).

### CD8^+^ T cell cross linking stimulation

For TCR stimulation naïve and primed CD8^+^ T cells were incubated for 20 min at 4°C with biotinylated anti-CD3ε mAb (5 and 1 μg/mL, BD Biosciences #553060). Cells were washed and stimulated for the indicated time by adding streptavidin (20 μg/mL, final concentration). For peptide stimulations purified primed CD8^+^ T cells were stimulated by hgp100_25–33_ peptide, KVPRNQDWL (AnaSpec, CliniSciences) at 1000 ng/mL.

### Western blotting

Stimulated cells were lysed at 4 °C for 10 min in 1% NP-40 lysis buffer (50 mM Tris pH 7.4, 150 mM NaCl, 5 mM EDTA, protease inhibitor cocktail (Roche #11836170001), 1 mM Na3VO4, 0.1% SDS). Samples were resolved by 10% SDS–polyacrylamide gel electrophoresis experiments. Blots were incubated overnight at 4 °C with the corresponding primary antibody directed p-AKT (Cell Signaling Technology #9271), Akt (Cell Signaling Technology #9272), p-Erk1/2 (Cell Signaling Technology #4377), Erk-1/2 (Cell Signaling Technology #9102) and β-Actin (Cell Signaling Technology #3700). Blots were incubated with corresponding peroxidase–conjugated secondary antibodies (Millipore #DC02L; #DC03L) for 1 hr at room temperature. ECL (enhanced chemiluminescence; SuperSignal West Pico and SuperSignal West Femto, Pierce) was used to visualize protein bands.

### Cytokine production

Primed mouse CD8^+^ T cells were cultured alone, or with B16 WT or B16-hgp100 target cells or stimulated with 200ng/mL PMA and 1μg/mL ionomycin (Sigma). After four hours of incubation at 37°C in the presence of FITC coupled anti-CD107a antibody (BD Pharmingen #553793), golgistop and golgiplug (BD Biosciences), cells were stained and the percentages of CD8^+^ T cells positive for CD107a, TNF-α (APC, Invitrogen, #17-7321-82) and IFN-γ (PE, Invitrogen, #12-7311-81) were measured by flow cytometry.

### Cytotoxicity

Target B16 cells were stained with 4μM of Cell Proliferation Dye eFluorTM 670 (Life Technologies) according to manufacturer’s instructions. Primed CD8^+^ T cells were then incubated with target cells for four hours at 37°C at different effector to target (E:T) ratios. Target cell killing was measured using CellEvent™ Caspase-3/7 Green Detection Reagent (Life Technologies) and analyzed by flow cytometry.

### Conjugate formation

This method has been adapted from a previous report on NK cells.^36^ Here, primed CD8^+^ T cells were incubated for 30 min on ice with CD8-APC-EF780 antibody (Invitrogen, #47-0081-82) in serum-free RPMI medium. They were then washed and resuspended at 20 × 10^6^ cells per ml. 100 μL of cell suspension was then added to 100 μL of labeled with Cell Trace Violet (V450) (Invitrogen) B16 WT or B16-hgp100 cells (at 20 × 10^6^ cells per mL) and centrifuged at 1,500 rpm (4 °C). After removing 150 μL of supernatant, cells were stimulated by incubation at 37 °C for 0, 5, or 10 min. Reactions were stopped by adding ice-cold phosphate-buffered saline. Conjugates were detected by flow cytometry as double positive CD8+ V450+ events.

### *In vivo* migration assays

Primed CD8^+^ T cells isolated from WT or *Dok1*/*Dok2* DKO mice were loaded with Cell Trace Violet Stain (Life Technologies #C34557) or Cell Trace Far Red DDAO (Life Technologies #C34553). Cells were mixed at a ratio of 1:1 and 10 × 10^6^ cells were *i*.*v*. injected in C57BL/6 mice. Then, 1 h later, recipient mice were euthanized, and blood, spleen, and lymph nodes were removed for quantification of Cell Trace Violet-labeled and Cell Trace Far Red-labeled T cells by flow cytometry.

### Adoptive cell transfer

For immunotherapy, Ly5.1 C57BL/6 (Janvier labs) were implanted with subcutaneous B16 melanoma (5×10^5^cells). At 10 days after tumor implantation, mice (n ≥ 5 for all groups) were sub-lethally irradiated (600 cGy), randomized, and injected intravenously with 5×10^5^ Pmel-1 DOK1, DOK2 KO or WT Ly5.2 primed CD8^+^ T cells and received intraperitoneal injections of IL-2 in PBS (6 × 10^4^ IU/0.5 mL) once daily for 3 days starting on the day of cell transfer. Mice with tumors greater than 400mm^2^ or in illness state were euthanized.

### Statistical Analysis

The data were expressed as the mean ± SEM. Prism 5.03 software (GraphPad, San Diego, CA, USA) was used for all statistical analysis. Statistical significance between control and DOK1 DOK2 KO groups was determined by two-tailed Student’s t-test. *P < 0.05, **P < 0.01, ***P < 0.001.

## RESULTS

### Primed DOK1 DOK2 KO CD8^+^ T lymphocytes have an effector memory phenotype

To understand how DOK1 and DOK2 regulate CD8^+^ T cell activity and especially their cytotoxic function against cancer cells, we crossed DOK-1/2 DKO (double KO) mice with pmel-1 TCR transgenic mice.^23^ We first tested the impact of DOK1 and DOK2 deletion on naïve and *in vitro* amplified CD8^+^ T cells. DOK-1/2 DKO and WT resting CD8^+^ T cells show similar proportions of CD4^+^ and CD8^+^ T cells, naïve (CD62L+CD44-), central memory (CD62L+CD44+) and effector memory (CD62L-CD44+) T cell subsets. To prime cells, naïve CD8^+^ T lymphocytes from spleen were purified and then expanded for 5 days, with anti-CD3, anti-CD28 or hgp-100 and IL-2 for 3 days followed by 2 days in IL-2 only (Fig. 1A). T cell subset phenotype was followed over the time at day 3 and day 5 by flow cytometry. Although no difference in proportion of Naïve (CD44-CD62L+), Central memory (CD44+CD62L+) and Effector memory (CD44+CD62L-) was observed in unstimulated CD8^+^ T cells or at day 3 of expansion, (Fig. 1B) we noticed that DOK1 DOK2 KO CD8^+^ T cells had a higher proportion of effector memory cells compared to WT cells at the day 5 of expansion (Fig. 1C). We identified the difference of CD62L expression in WT and DOK-1/2 DKO CD8^+^ T cells between day 3 and day 5. (Fig. 1D) Therefore, DOK1 and DOK2 regulate the formation of memory CD8^+^ T cells.

**Figure 1.**
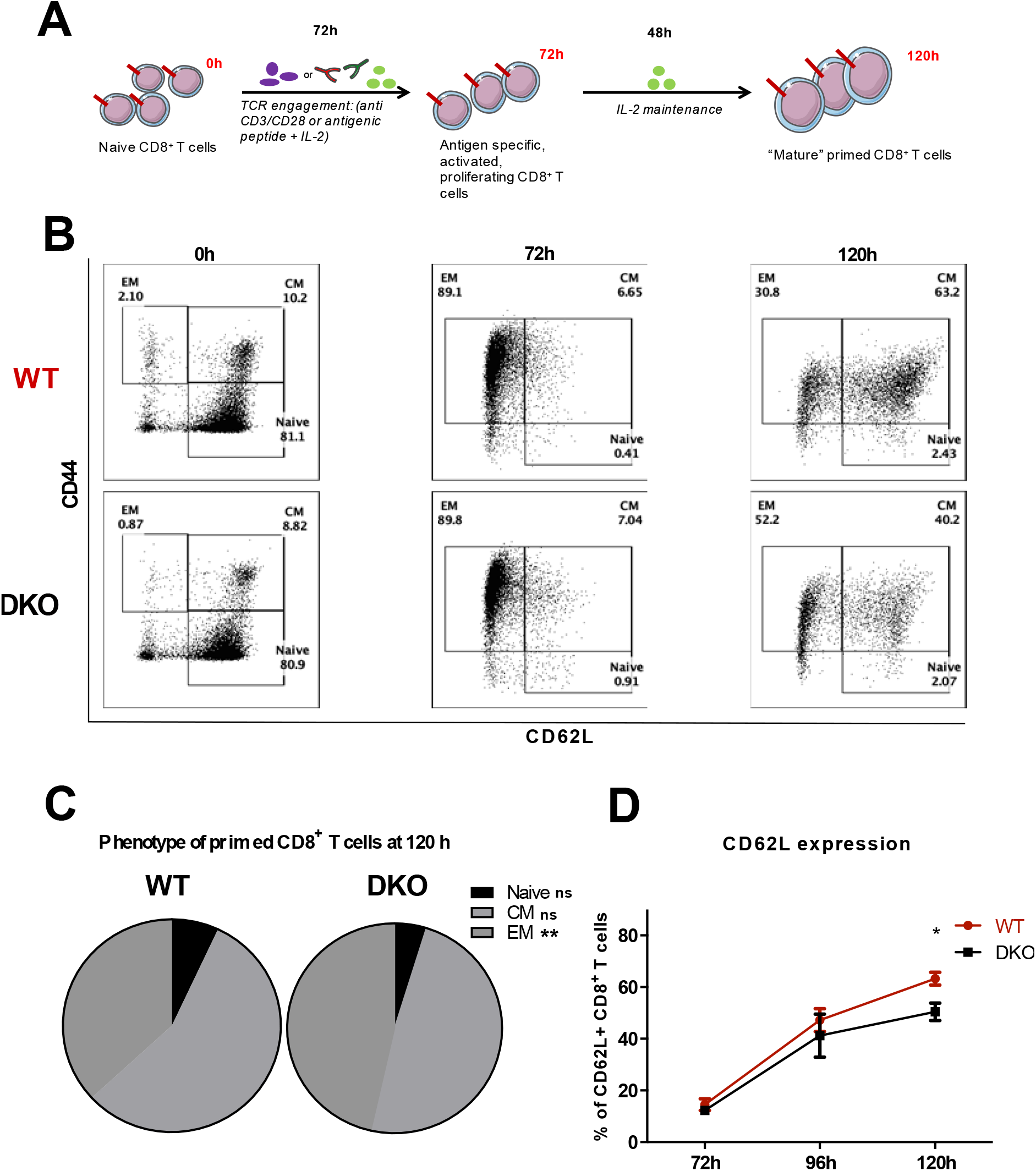
Dok1 Dok2 KO primed CD8^+^ T cells have effector memory phenotype. A) Used CD8^+^ T cell expansion protocol: 3 days of TCR engagement with anti-CD3 2µg/ml and CD28 1μg/ml or hgp-100 peptide (100ng/ml) in the presence of IL-2 100 UI/ml; following by 2 days of maintenance on IL-2 (100UI/ml) B) Representative histogram showing the phenotype of primed CD8^+^ T cells at 0h, 72h and 120h of expansion (n=12). C) Effector memory (EM: CD44+CD62L-), Central memory (CM: CD44+ CD62L+) and Naïve (CD44-CD62L+) proportion in primed CD8^+^ T cells at 120h of expansion (n=15). D) Expression of CD62L during expansion of CD8+ T cells measured by flow cytometry (n=3). Error bars, SEM. *, p<0,05; **, p<0,01 by Student t-test.

### DOK1 and DOK2 invalidation improves TCR signaling in primed CD8^+^ T cells

We, then, sought to explore the role of DOK1 and DOK2 invalidation in TCR signaling. We used two doses (5 and 1 μg/mL) of biotinylated anti-CD3 to determine the optimal dose for CD8^+^ T cells stimulation (Fig. S1A). *Dok1*/*Dok2* DKO and WT CD8^+^ T cells purified from spleen were stimulated with biotinylated anti-CD3 and cross-linked with streptavidin during the indicated time. After cell lysis the levels of pErk and pAkt were evaluated by Western blot (WB) analysis. Naïve WT and DKO CD8^+^ T cells show the same level of pErk and pAkt upon TCR stimulation (Fig. 2A).

**Figure 2.**
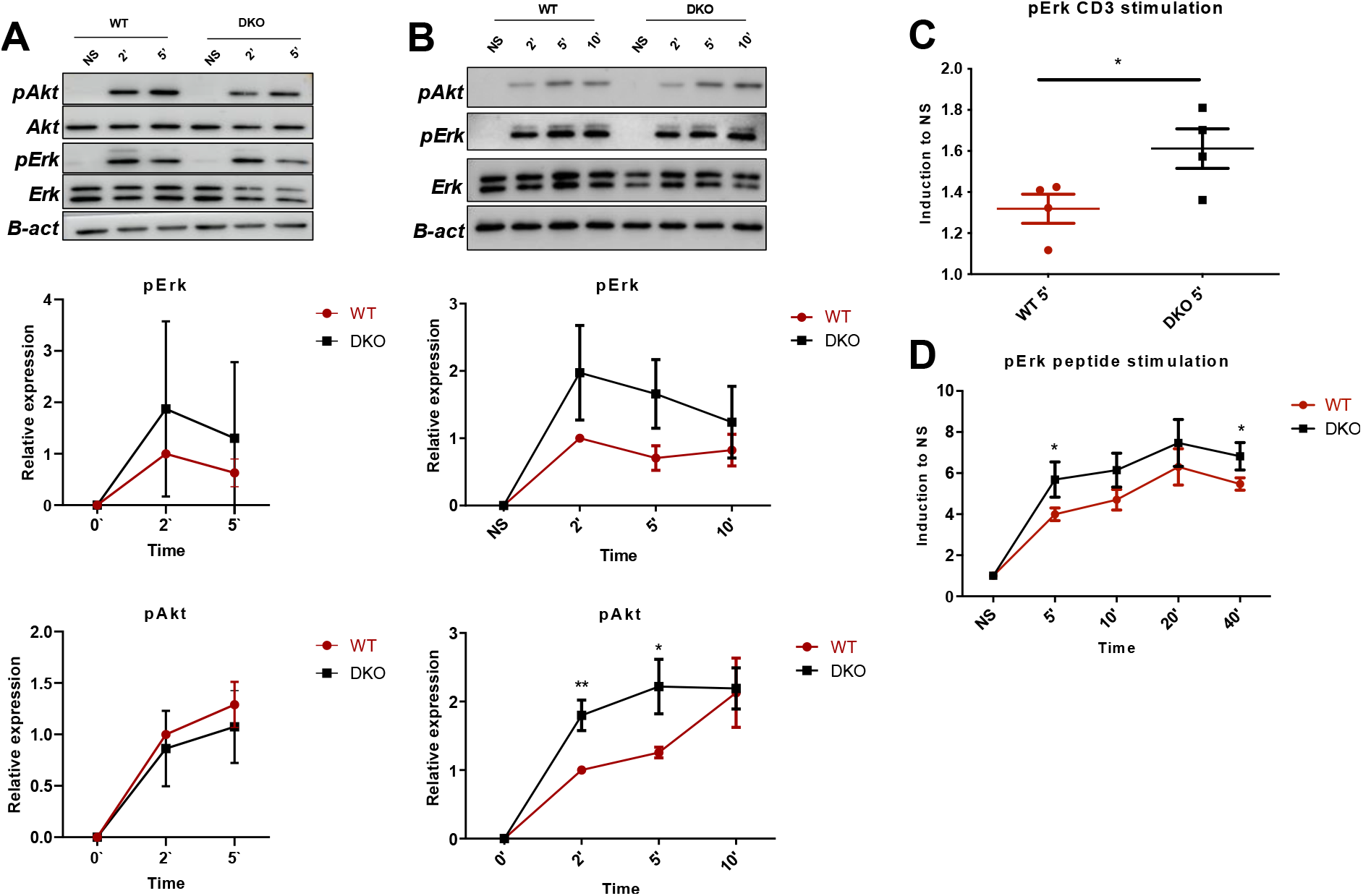
Dok1 Dok2 KO improves TCR signaling in primed CD8+ T cells. Naïve and primed CD8+ T cells were stimulated by anti-CD3 5µg/ml. A) Representative immunoblots of naïve cells stimulated with anti-CD3 for 2 and 5 minutes. Normalized quantification of pErk and pAkt induction is shown. (n=3) B) Representative immunoblots of primed CD8+ T cells stimulated with anti-CD3 for 2, 5 and 10 minutes. Normalized quantification of pErk and pAkt induction is shown. (n=4). C) pErk induction in primed CD8+ T cells after 5 minutes of stimulation by anti-CD3. Phopho-Flow detection method. (n=4). D) pErk induction in primed CD8+ T cells after TCR stimulation with hgp-100 peptide 1µg/ml for 5, 10, 20 and 40 minutes. Phopho-Flow detection method. (n=5). E) Representative histogram of pErk induction in WT and DKO primed CD8+ T cells after 40 minutes of stimulation by hgp100 peptide 1µg/ml. Error bars, SEM. *, p<0,05,**, p<0,01 by Student t-test.

To determine whether the loss of DOK1 and DOK2 affects TCR signaling in primed CD8^+^ T cells, similar experiments were performed. Primed *Dok1*/*Dok2* DKO CD8^+^ lymphocytes showed an upregulation of pErk and pAkt expression compared to WT CD8^+^ T cells upon TCR stimulation, although only pAkt appeared to be statistically significant (Fig. 2B). Subsequently, phosphoflow experiments were performed. Primed CD8^+^ T cells were stimulated with biotinylated anti-CD3 and cross-linked with streptavidin during 2, 5 and 10 minutes. Phosphorylation of ERK1/2 was detected by flow cytometry (Fig. 2C). Again, pErk expression was increased in primed *Dok1*/*Dok2* DKO CD8^+^ T cells compared to WT cells, confirming our WB experiments.

To ensure that this TCR signaling improvement in *Dok1*/*Dok2* DKO primed CD8^+^ T lymphocytes is conserved with a different TCR stimulation setting, we performed a stimulation with hgp-100 peptide. This peptide is specifically recognized the transgenic TCR expressed at the surface of pmel-1^+^ CD8^+^ T cells. Cells were peptide-stimulated, immediately fixed and stained with antibodies against pErk and pS6. Flow cytometry analysis revealed an increase of pErk induction in *Dok1*/*Dok2* KO primed CD8^+^ T lymphocytes compared to WT CD8^+^ T cells (Fig. 2D). No difference in pS6 expression was found (data not shown).

Similarly, we performed WB analysis of primed CD8^+^ T cells lysates after a stimulation with hgp-100 peptide. We didn’t notice any difference in pErk and pAkt expression levels between WT and DKO cells. (Fig. S2A-C)

Altogether, these findings suggest that DOK1 and DOK2 deficiency enhance TCR signaling upon different stimulation settings. This effect was only observed when CD8^+^ T cells were primed *in vitro*.

### Dok1/Dok2 DKO and WT CD8^+^ T cells show similar cytotoxicity in vitro

To assess CD8^+^ T cell cytotoxicity a murine B16 melanoma cell line expressing constitutively hgp-100 antigen was used. Primed pmel-1^+^ CD8^+^ T cells can recognize hgp-100 antigen at the surface of B16-hgp-100 expressing cells but not when the peptide is not expressed. Primed *Dok1*/*Dok2* DKO and WT CD8^+^ T cells were co-cultured with B16-hgp-100 cells at indicated effector/Target (E/T) ratio. Expression of IFN-γ and TNF-α was detected by flow cytometry (Fig. S3A). Surprisingly, *Dok1*/*Dok2* DKO and WT CD8^+^ T cells expressed the same level of IFN-γ and TNF-α at any tested E/T ratios. (Fig. 3A and data not shown) Likewise, degranulation marker CD107a showed also similar expression between *Dok1*/*Dok2* DKO and WT CD8^+^ T cells (Fig. 3B).

**Figure 3.**
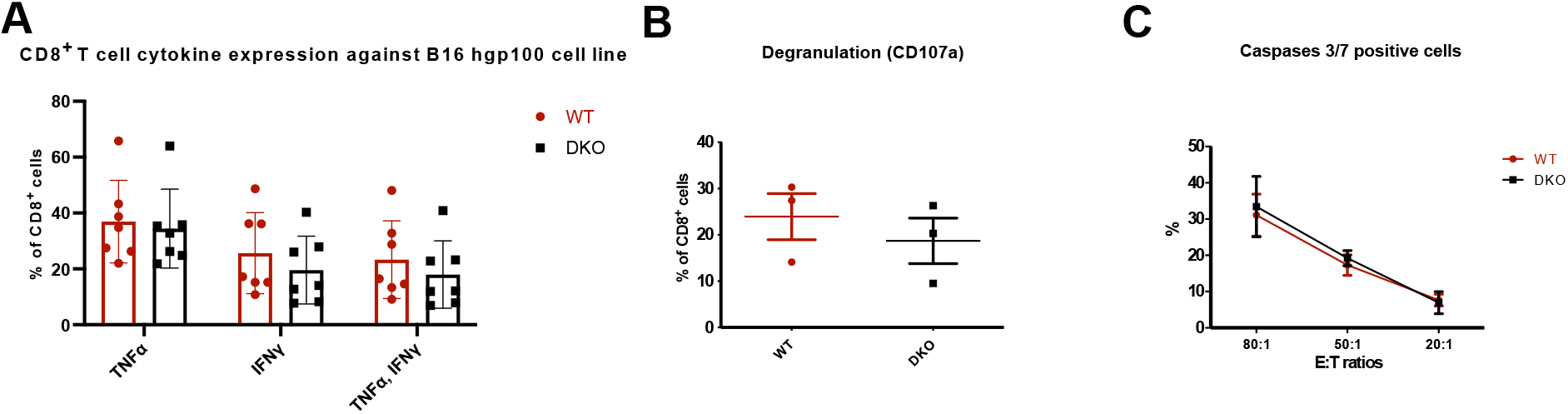
Dok1 Dok2 KO does not affect cytotoxic function of primed CD8+ T cells. A) Cytokine production by primed CD8+ T cells at 1:10 E:T ratio after 4h of co-culture with B16 hgp100 target cells measured by flow cytometry (n=7) B) Degranulation measured by CD107a expression by primed CD8+ T cells at 1:10 E:T ratio after 4h of co-culture with B16 hgp100 target cells measured by flow cytometry (n=3) C) Caspases 3/7 activation in B16 hgp100 cells after co-culture with primed CD8+ T cells in different E:T ratios measured by flow cytometry (n=3)

Next, the capacity of primed CD8^+^ T cells to kill B16-hgp100 cells was analyzed by measuring caspase 3/7 activation in target cells after 4h of co-culture. At any tested ratios WT and DOK1 DOK2 KO primed CD8^+^ T cells showed the same cytotoxic activity (Fig. 3C) Therefore, these data suggest that DOK1 DOK2 KO and WT CD8^+^ T cells show similar cytotoxicity *in vitro*.

### DOK1 and DOK2 KO primed CD8^+^ T cells do not improve survival of tumor bearing mice

Finally, we tested the capacity of primed CD8^+^ T cells to rescue the development of tumor after adoptive cell transfer. We made a subcutaneous B16-hgp-100 tumor injection and 10 days after we performed an adoptive cell transfer of WT and DOK1 DOK2 KO CD8^+^ T cells. PBS injection was used in a control group. We found that adoptive cell transfer improved a survival of tumor bearing mice, however, there was no difference between WT and DOK1 DOK2 KO group. (Fig. S4). Thus, this data confirms the *in vitro* experiments and DOK1 DOK2 invalidation do not improve primed CD8^+^ T cell cytotoxicity in the context of adoptive cell transfer.

### DOK1 DOK2 KO and WT CD8^+^ T cells show similar effector: target conjugate formation and migration

Since, *Dok1*/*Dok2* DKO primed CD8^+^ T lymphocytes have fewer expression of CD62L adhesion molecule (Fig. 1D), we hypothesized that it could impact the formation of effector-target cell conjugates or migration. For migration experiment WT and *Dok1*/*Dok2* DKO primed CD8^+^ T cells were labeled with dyes of different color. Next cells were mixed in proportion 1:1 and 1×10^6^ of cells were intravenously injected in healthy mice. One hour after injection blood, spleen and lymph nodes were taken to evaluate the proportion of WT and DOK-1/2 DKO CD8^+^ T in each organ. No difference in cell proportion was detected (Fig. S5 A-D). To evaluate conjugate formation, we stained target cells with Cell Trace Violet and effector cells with fluorochrome-coupled CD8 mAb. A co-culture experiment was performed, and conjugate formation was followed during time course (Fig. S5E). WT and *Dok1/Dok2* DKO primed CD8^+^ T cells showed the same effector-target conjugate formation.

## DISCUSSION

In this study, we examined the role of DOK1 and DOK2 in naïve and primed CD8^+^ T cells. Primed but not naïve *Dok1*/*Dok2* DKO CD8^+^ T cells had an improved TCR signaling and showed more effector memory subtypes compared to WT CD8^+^ T cells. Building on these results we hypothesized that DOK1 and DOK2 invalidation in CD8^+^ T cells may be a promising approach to improve their anti-tumor functions and thus subsequent immunotherapy. However, the phenotypic and signaling differences did not translate into a difference in cytotoxic response.

In the present study we showed that DOK1 and DOK2 regulate *in vitro* memory CD8^+^ T cell formation. We found that at the end of 5 days of CD8^+^ T cell expansion we have more effector memory phenotype in *Dok1*/*Dok2* DKO CD8^+^ T cells. We believe that this is due to a slower re-expression of CD62L between day 3 and day 5 when TCR stimulation was canceled. We think that during this period, cells are only proliferating due to IL-2 presence, but T cells subset are starting to “rest” from TCR previous stimulation. At least in CD8^+^ T cells, DOK1 and DOK2 seem to exert their inhibiting role to favorize the activated cells to go back to a “resting” state, but we could not detect any difference at naïve state, not only in signaling but also in phenotypic experiments.

CD62L or L-selectin controls T-cell migration and is negatively controlled by PI3K-Akt pathway activation.^37^ Previously it was shown that DOK1 negatively controls SDF-1α induced cell migration.^38^ Thus, we performed *in vivo* migration experiments using primed CD8^+^ T cells but we could not find the difference between WT and *Dok1*/*Dok2* DKO T cells.

Previous studies showed major improvements of TCR signaling, proliferation and cytokine production in naïve and memory DOK1 DOK2 KO CD4^+^ T cells.^29,39^ Only few studies were performed on DOK1 DOK2 invalidation in CD8^+^ T cells.^31,40^ In agreement with our results WT and DOK1 DOK2 KO naïve CD8^+^ T cells showed similar signaling upon TCR stimulation. Therefore, we confirm the difference of DOK1 and DOK2 regulation of TCR signaling in CD4^+^ and CD8^+^ naïve T cells. Considering fundamental differences in the role of CD4^+^ and CD8^+^ T cells and their complex functional interplay, it is rational to suggest some differences in TCR signalosome of these cells.^41–43^ For example, recently a crucial difference in TCR initiation signaling was revealed showing that LCK binding is stronger to CD8 compared to CD4 coreceptor, leading to more potent intracellular signaling.^44^ We found that primed *Dok1*/*Dok2* DKO CD8^+^ T cells showed an improved TCR signaling by upregulation of pAkt and pErk upon TCR engagement with two different methods of stimulation (agonist CD3 mAb and peptide) and two kind of experimental approaches. Naïve and memory T cells have considerable differences in their function physiology and TCR signalosome. Particularly, it was demonstrated that memory CD8^+^ T cells have more CD8-bound Lck than naïve cells and CD4^+^ T cells have less Zap70 and Slp76 phosphorylation upon TCR stimulation, suggesting a faster and more efficient signal transduction pathway in memory T cells.^19,20^ These data confirm our findings and suggest some functional or structural differences in CD8^+^ naïve versus primed signalosomes. Further investigation notably using high throughput technologies such as mass spectrometry are needed to decipher these phenomenon.^13^

Compensation mechanisms and signaling re-wiring may also occur in TCR signaling pathway, like the inhibitory TCR signaling protein Csk, normally associated with PAG, could associate with another protein PTPN22 in absence of PAG compensating TCR signaling.^39^ DOK1 and DOK2 could be seen as a platform to recruit other inhibitory proteins (RasGAP, SHIP, Csk). Maybe other proteins could compensate the lack of DOK1 and DOK2 when these proteins are totally absent. Thus, methods to downregulate transiently DOK1 DOK2 in CD8^+^ T lymphocytes by shRNA or CRISPRi-Cas9 techniques may be interesting to avoid these compensation mechanisms and understand more precisely DOK1 and DOK2 regulation.

In this study we wanted to assess the capacity of TCR signaling inhibitory proteins DOK1 and DOK2 to improve CD8^+^ T cells immunotherapy. In the development of T cell-based immunotherapy the problem of T cell functionality blunting in the tumor microenvironment is crucial. The concept of improving the strength of TCR signaling upon TCR activation is very important to overcome this problem. Therefore, we tested TCR signal inhibiting proteins DOK1 and DOK2 as potential candidates to increase TCR signaling in CD8^+^ T cells. The fact that we found the increase of pErk and pAkt in primed but not naïve CD8^+^ T cells is even advantageous in this context, as only primed CD8^+^ T cells are used for adoptive cell transfer immunotherapies nowadays. Previously, it was shown that inhibition of Akt pathway by rapamycin could improve the generation of memory cells in terms of their quantity and quality.^45^ The acquisition of effector functions of CD8^+^ T cells associated with intense Akt signaling impairs the *in vivo* antitumor efficacy of adoptively transferred cells.^46^ The emerging consensus on this question is that central memory tumor-reactive CD8^+^ T cells have an improved antitumor capacity in comparison with effector memory cells.^47,48^ Therefore, the activation both pErk and pAkt as we can see in the context of DOK1 DOK2 invalidation would be not advantageous for antitumor capacity of primed CD8^+^ T cells as the positive influence of pErk upregulation would be compensated by the negative influence of effector memory phenotype due to increased pAkt. Probably in the concept of CD8^+^ T cells immunotherapy improvement by acting through TCR signaling inhibiting such polyvalent inhibiting proteins as DOK1 and DOK2 would be excessive, and proteins acting on inhibiting of one specific signaling pathway would fit more. As successful examples of targeting intracellular inhibiting proteins in the context of cancer immunotherapy, we can mention recently adapted for clinical trials CISH, targeting PLC-γ1 and HPK-1, targeting SLP-76.^23,49^ Both could be associated to Erk pathway improvement, without direct effect on Akt pathway.

In summary, our data provided evidence that DOK1 and DOK2 interfere in primed CD8^+^ T cell TCR signaling negative regulation and have impact on memory CD8^+^ T cell formation. We underlined an interesting phenomenon that DOK1 and DOK2 could play a different role in naïve and memory TCR signaling, however based on our model the DOK1/DOK2 adaptor proteins do not appear to be good candidates for CD8^+^ T cell manipulation in immuno-oncology. Therefore, due to complexity of TCR signaling there is a real need of screening studies of invalidation of TCR signaling inhibitory proteins to improve existing CD8^+^ T cell-based immunotherapies.

## ACKNOWLEDGEMENTS

This work is supported by institutional grants from the Institut National de la Santé et de la Recherche Médicale (Inserm), Centre National de la Recherche Scientifique (CNRS) and Aix-Marseille Université to CRCM; by project grants from the Fondation ARC pour la recherche sur le Cancer (PJA20161204835), Groupement des Entreprises Françaises dans la Lutte contre le Cancer (GEFLUC) Marseille-Provence. The “Immunity and Cancer” team was labeled by the Fondation pour la Recherche Médicale “Equipe FRM DEQ20180339209”. D.O. is Senior Scholar of the Institut Universitaire de France. V.L. was supported by a doctoral fellowship from the Ministère de l’Education et la Recherche, then by the Fondation ARC pour la recherche sur le Cancer. P-L.B. is supported by a doctoral fellowship from Ligue Nationale contre le Cancer and. G.G. was supported by a post-doctoral fellowship from the Fondation ARC pour la recherche sur le Cancer, then by the Janssen Horizon Fonds de dotation. We are grateful to the staff of the CRCM animal facility for taking care of the mouse strain colonies and the CRCM cytometry platform for FACS analysis.

## AUTHOR CONTRIBUTIONS

V.L., P-L.B., C.M., R.C. and G.G. all performed experiments. V.L., G.G. and J.A.N. designed experiments and analyzed the data. Y.Y. provided a crucial mouse model for this study. D.O. contributed to some funding acquisitions. G.G. and J.A.N. led the research program. V.L., G.G., and J.A.N. wrote the manuscript.

## CONFLICTS OF INTEREST

D.O. is a cofounder of Imcheck Therapeutics, Alderaan Biotechnology and Emergence Therapeutics. The other authors declare no competing financial interests.

