## supplemental figures and legends for "DOK1 and DOK2 regulate CD8^+^ T cell signaling and memory formation without affecting tumor cell killing"

### Supplementary Figures

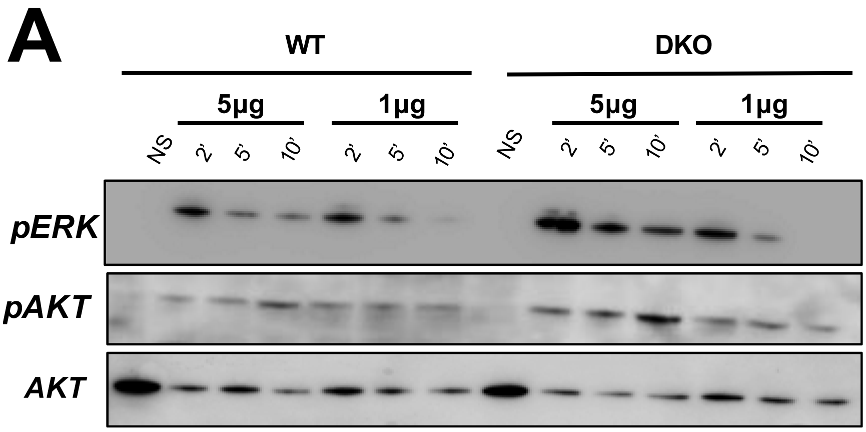


**Supplementary Figure 1. Validation of WB TCR signaling in primed CD8+ T cells.** A) Primed CD8+ T cells were stimulated by anti-CD3 5µg/ml and 1µg/ml. Representative immunoblot of 3 experiments is shown.

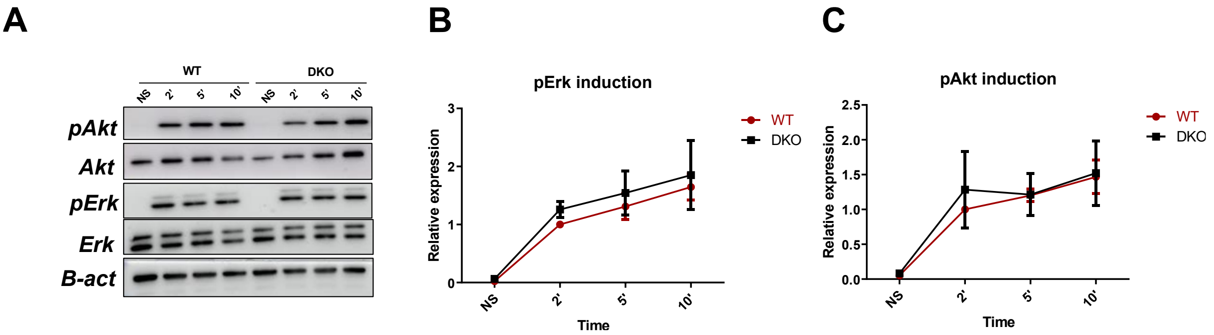


**Supplementary Figure 2. Dok1 Dok2 KO does not improve TCR signaling in primed CD8+ T cells after peptide stimulation.** A) Primed CD8+ T cells were stimulated by hgp-100 peptide 1000ng/ml. Representative immunoblot of primed cells stimulated with hgp-100 for 2, 5 and 10 minutes. Normalized quantification of pErk (B) and pAkt (C) induction is shown (n=4).

**A**

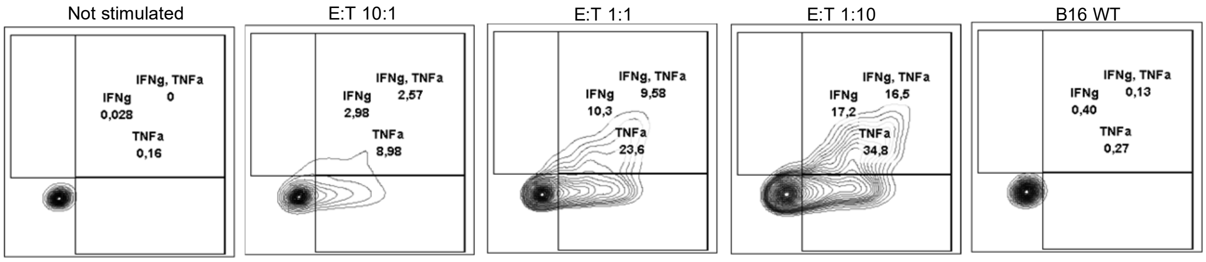


**Supplementary Figure 3. Validation of cytotoxicity against B16$^{hgp100}$ cells line.** Histograms showing cytokine production of unstimulated primed WT CD8+ T cells after 4h of stimulation with B16-WT or B16 expressing hgp100 in different E:T ratios.

A

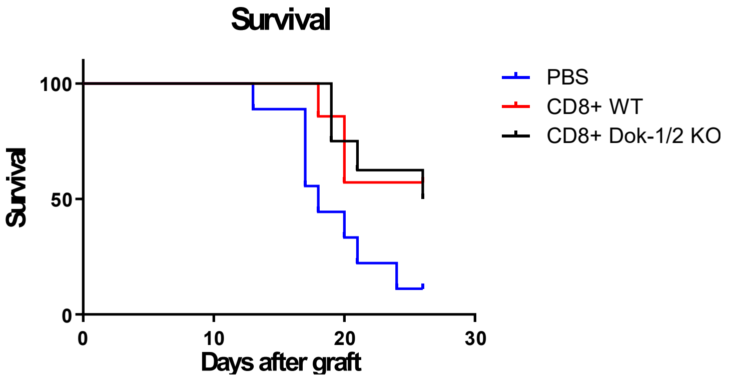


**Supplementary Figure 4. DOK1 DOK2 KO adoptive primed CD8+ T cell transfer do not improve survival in cancer model** A) Mouse survival after B16 hgp100 implantation and treatment with PBS, WT or DOK1 DOK2 KO primed CD8+ T cells (n>=5).

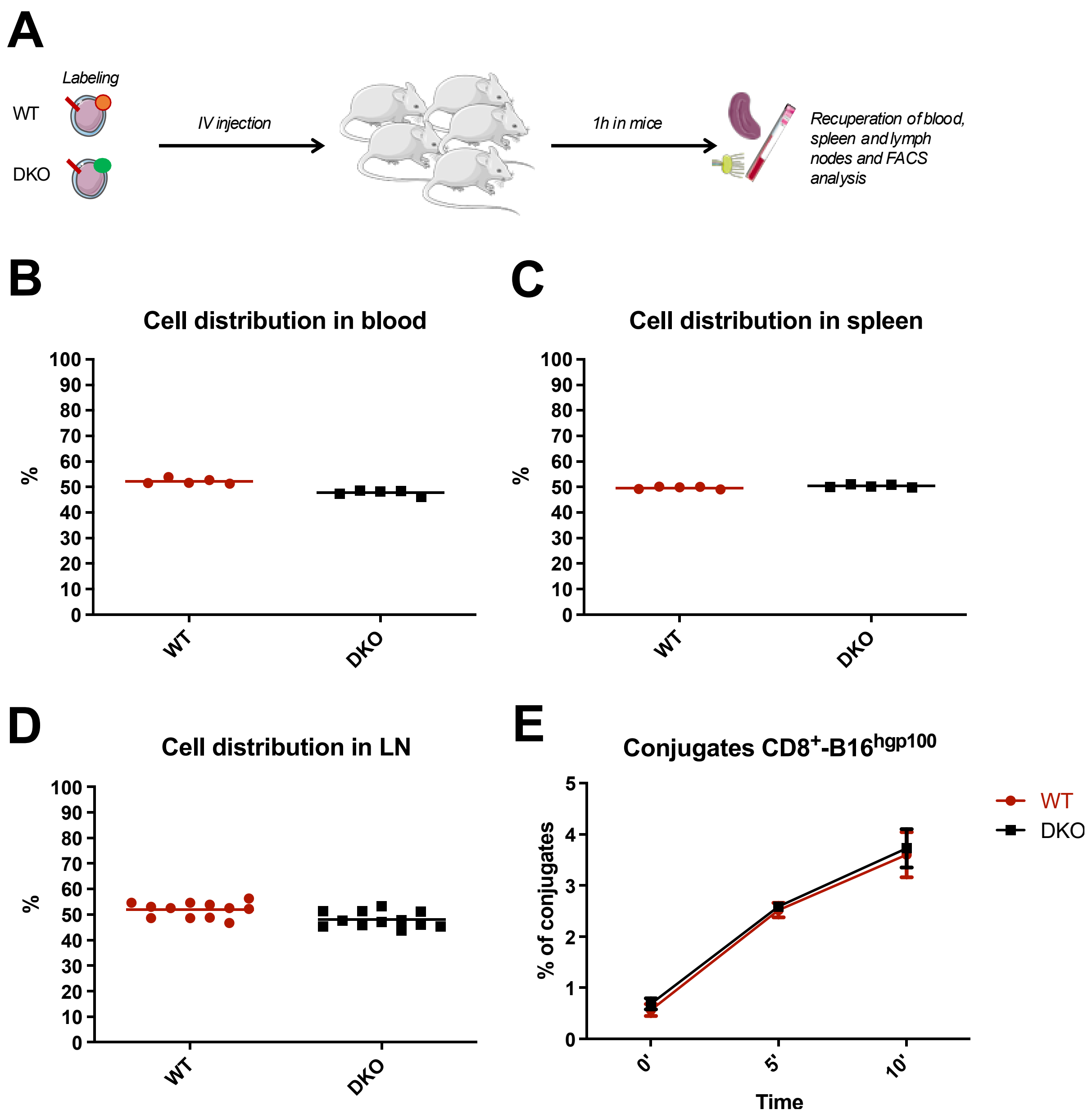


**Supplementary Figure 5. Dok1 Dok2 KO does not affect primed CD8$^+$ T cells *in vivo* migration and cell conjugate formation.** A) Experimental protocol of *in vivo* primed CD8$^+$ T cell migration. WT and DKO cell distribution in blood (B), spleen (C) and lymph nodes (D) is shown (n=5). E) Conjugate formation between primed CD8$^+$ T cells and B16 hgp100 cells at 1:1 ratio after contact and co-culture for 5 and 10 minutes measured by flow cytometry (n=4).
